# The “glue-gun ant” *Lordomyrma azumai* uses an adhesive thread secretion to immobilize opponents

**DOI:** 10.64898/2026.10.07.757171

**Authors:** Hajime Sasaki, Riou Mizuno, Kenshin Sato, Hayato Shinohara, Adam Khalife, Karen Kohama, Fuminori Ito

## Abstract

In stinging Hymenoptera, the poison gland and its vesicle have been understood largely as sources of toxic, irritating, or antimicrobial compounds, which the sting injects into a target^1,2^. However, external uses of poison gland secretions have also been reported: *Pseudoneoponera* (formerly *Pachycondyla*) froth a proteinaceous poison gland secretion into foam threads that entangle attacking ants^3^, and *Crematogaster* smear opponents with a contact toxin generated on a spatulate sting from poison and Dufour gland products^4^. Here we report that workers of the myrmicine ant *Lordomyrma azumai* produce, in the poison vesicle, a secretion, likely proteinaceous, that solidifies upon extrusion into an adhesive thread. The sting is retained as the conduit, but the secretion it delivers is applied externally rather than injected, and acts as a structural material: workers wipe the thread onto opponents and tether them to the substrate. To our knowledge, no hymenopteran venom secretion functions as a solid thread; in the closest case, *Pseudoneoponera*^3^, the secretion entangles attackers as a foam rather than as a continuous solid filament.

---

When confronting heterospecific ants, *L. azumai* workers orient the gaster toward the opponent and extrude an adhesive thread from the sting, actively wiping it onto the opponent’s cuticle and the surrounding soil (Figure 1A; Video S1). Viscous within the poison vesicle, the secretion solidifies almost instantly after extrusion into a continuous filament (mean ± SD = 12.5 ± 4.6 µm in width, n = 14 measurements along a single thread; Figure 1B) that adheres to the opponent. Its deployment is context-dependent. In staged encounters within the nest, workers used thread attacks far more often than biting against five of the six sympatric competitor species tested (paired t-tests with colony as the unit of replication, n = 3 colonies; Figure S1A). The small, slow-moving *Vollenhovia emeryi* elicited mainly biting, with no significant difference between attack types. Prey mealworms were attacked only with mandibles, and non-nestmate conspecifics from both the same and different localities elicited no aggression at all. Outside the nest, solitary workers deployed the thread against *Nylanderia flavipes* and *Pheidole fervida* at similar frequencies (mean ± SD = 5.00 ± 3.56 and 5.60 ± 3.60 attacks per trial, n = 10 each; Wilcoxon rank-sum test, *W* = 41, *p* = 0.517) but adopted defensive postures (death feigning with retracted antennae, as previously noted in this species^5^) significantly more often against *P. fervida* than against *N. flavipes* (3.50 ± 5.52 versus 0.20 ± 0.42 postures per trial; *W* = 8.0, *p* < 0.001). At the nest entrance, workers combined gaster-based phragmosis, a form of nest-plugging usually achieved with the head in ants^6^, with thread attacks (Figure S1B). Intruders entered 19 of 30 empty control chambers, but only 6 of 30 nests (Figure S1B, C). All six entries occurred in a single colony and coincided with an absence of active defense, although defense was not experimentally manipulated.

**Figure 1.**
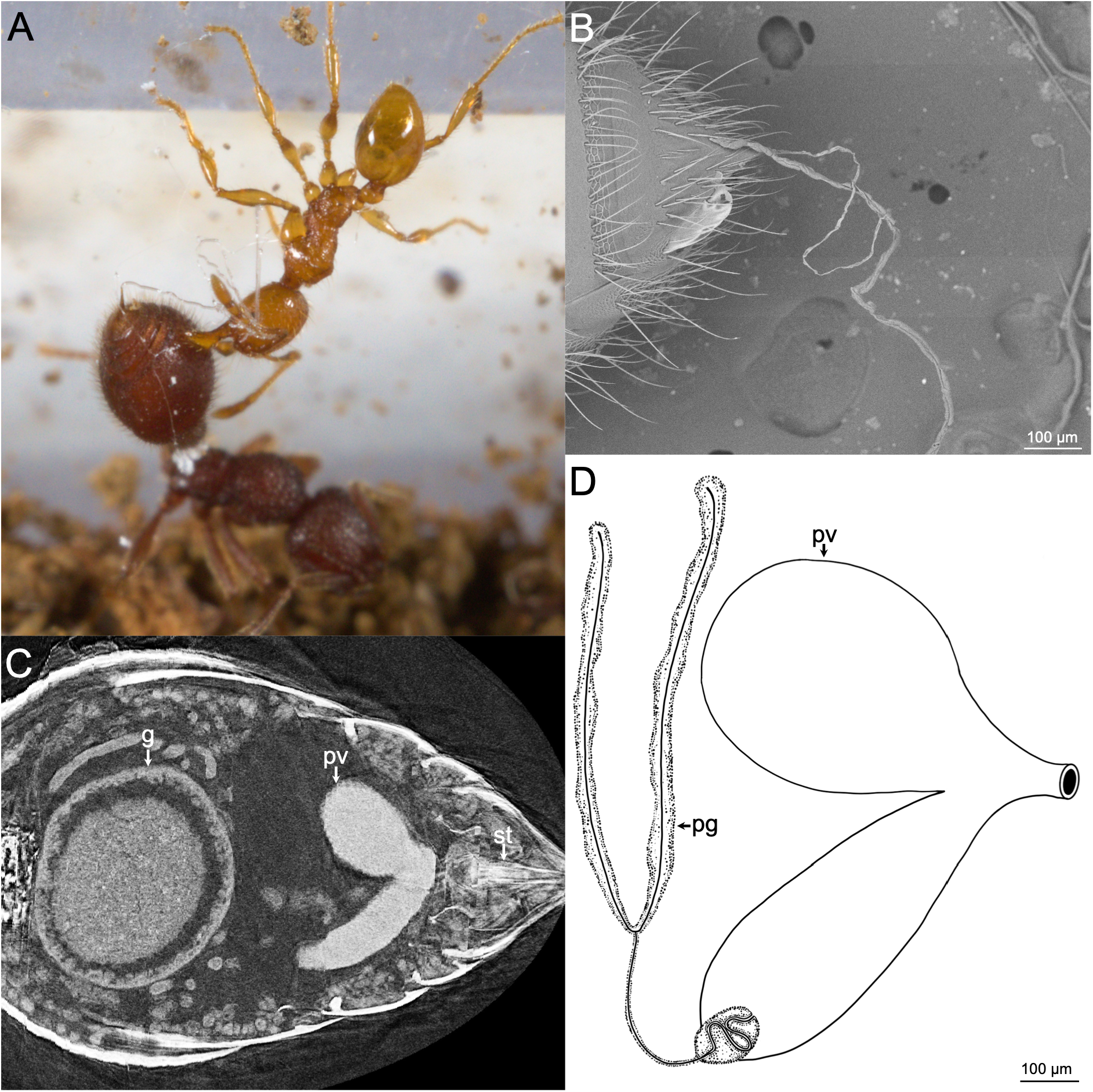
A venom-derived adhesive thread and the venom apparatus of *Lordomyrma azumai*. (A) A *L. azumai* worker (bottom) performing a sticky thread attack on a *Pheidole fervida* (top), with the adhesive thread visible between the *L. azumai* gaster and the opponent’s body. Scanning electron micrograph showing the adhesive thread secreted from the sting. (C) Dorsal-view micro-CT reconstruction of the gaster. (D) Illustration of the *L. azumai* worker poison gland and poison vesicle. Abbreviations: g, gut; pg, poison gland; pv, poison vesicle; st, sting.

The thread’s immobilizing power is amplified by environmental material. Immobilization success rates were significantly higher when soil was present compared to bare plaster floors (generalized linear mixed model with colony as random intercept; *N. flavipes*: OR = 6.54, 95% CI: 3.18-13.44, χ^2^ = 32.41, df = 1, p < 0.001; *P. fervida*: OR = 9.59, 95% CI: 4.79-19.20, χ^2^ = 53.11, df = 1, p < 0.001; Figure S1D). Soil particles adhered to the thread and tethered opponents directly to the surrounding soil. Immobilized opponents were then dismembered by nestmates; of nine tethered opponents monitored after attack, eight were dead within 24 hours. We propose the term “environmentally-assisted defense” for this synergy between a self-produced secretion and environmental material. Although functionally distinct, the incorporation of soil particles into defensive threads recalls the use of wood-reinforced silk by the distantly related ants *Melissotarsus* and *Rhopalomastix*^7^, illustrating convergent exploitation of environmental materials to augment self-produced secretions.

Dissection localized the viscous secretion to the poison vesicle, and micro-computed tomography revealed a prominent, bilobed vesicle occupying much of the posterior gaster (Figure 1C, D); the Dufour gland contained no comparable material. Scanning electron microscopy showed the secretion emerging directly from the sting tip as a single continuous filament (Figure 1B). Infrared spectroscopy showed broad Amide I and Amide II bands centered near 1645 and 1540 cm^−1^, consistent with a proteinaceous composition. Because residual water-vapor absorption precluded band-narrowing analysis of the Amide I contour^8^, we do not assign its component secondary structures. The full infrared spectra are available at Zenodo^9^.

Two features make this system notable. First, unlike conventional venom, the secretion is effective only after leaving the sting and solidifying. Second, the defense does not rely on the secretion alone; it co-opts soil from the nest floor. This raises the mean proportion of opponents immobilized from ∼9% to ∼38% for *N. flavipes* and from ∼10% to ∼50% for *P. fervida* compared with bare surfaces, amplifying a limited secretion. The thread is not used against prey, but appears to be reserved for heterospecific threats, suggesting that the secretion is both limited and deployed under tight behavioral control. Preliminary observations of similar behavior in an undescribed Malaysian *Lordomyrma* species suggest that the trait may not be confined to *L. azumai* (H. Sasaki and F.I., personal observation).

*Lordomyrma azumai*, and potentially other species in the genus *Lordomyrma*, thus offers a tractable system for studying how the poison gland, ancestrally a chemical weapon, can be evolutionarily repurposed into a physical one. In this ant, venom becomes glue, and soil becomes an anchor that pins the opponent down. More broadly, the system illustrates how defensive innovations can emerge through the functional transformation of ancestral traits rather than the evolution of entirely new structures.

## ACKNOWLEDGMENTS

We thank Toshiharu Akino (Kyoto Institute of Technology) for his kind suggestions regarding the chemical analysis of the venom. We acknowledge Shunsuke Izumi (Hiroshima University) for helpful discussions on the chemical analysis of the secretion. We are grateful to Shogo Makita and Hiroki Matsumura (Kagawa University) for collecting and providing ant colonies; Hiroki Matsumura also provided valuable suggestions. We thank Evan P. Economo (Okinawa Institute of Science and Technology Graduate University, OIST) and the OIST Imaging Section, especially Shinya Komoto, for providing access to the Zeiss Xradia micro-CT scanner. This work was supported by the Fujiwara Natural History Foundation (awarded to H. Sasaki).

## SUPPLEMENTAL INFORMATION

Supplemental Information includes experimental procedures, one figure, and one video, and can be found with this article online at *bxs.

Document S1. Experimental Procedures and One Figure.

Video S1. Context-dependent sticky-thread attack and environmentally-assisted immobilization by *Lordomyrma azumai*. The single file comprises three labeled segments: (1) a worker secretes an adhesive thread from the sting and wipes it onto a heterospecific opponent; (2) the thread adheres to the opponent and entangles soil grains, tethering it to the substrate; (3) the viscous secretion emerges from the sting and rapidly solidifies into a continuous filament.

