## Supplementary material for "The “glue-gun ant” *Lordomyrma azumai* uses an adhesive thread secretion to immobilize opponents": Document S1

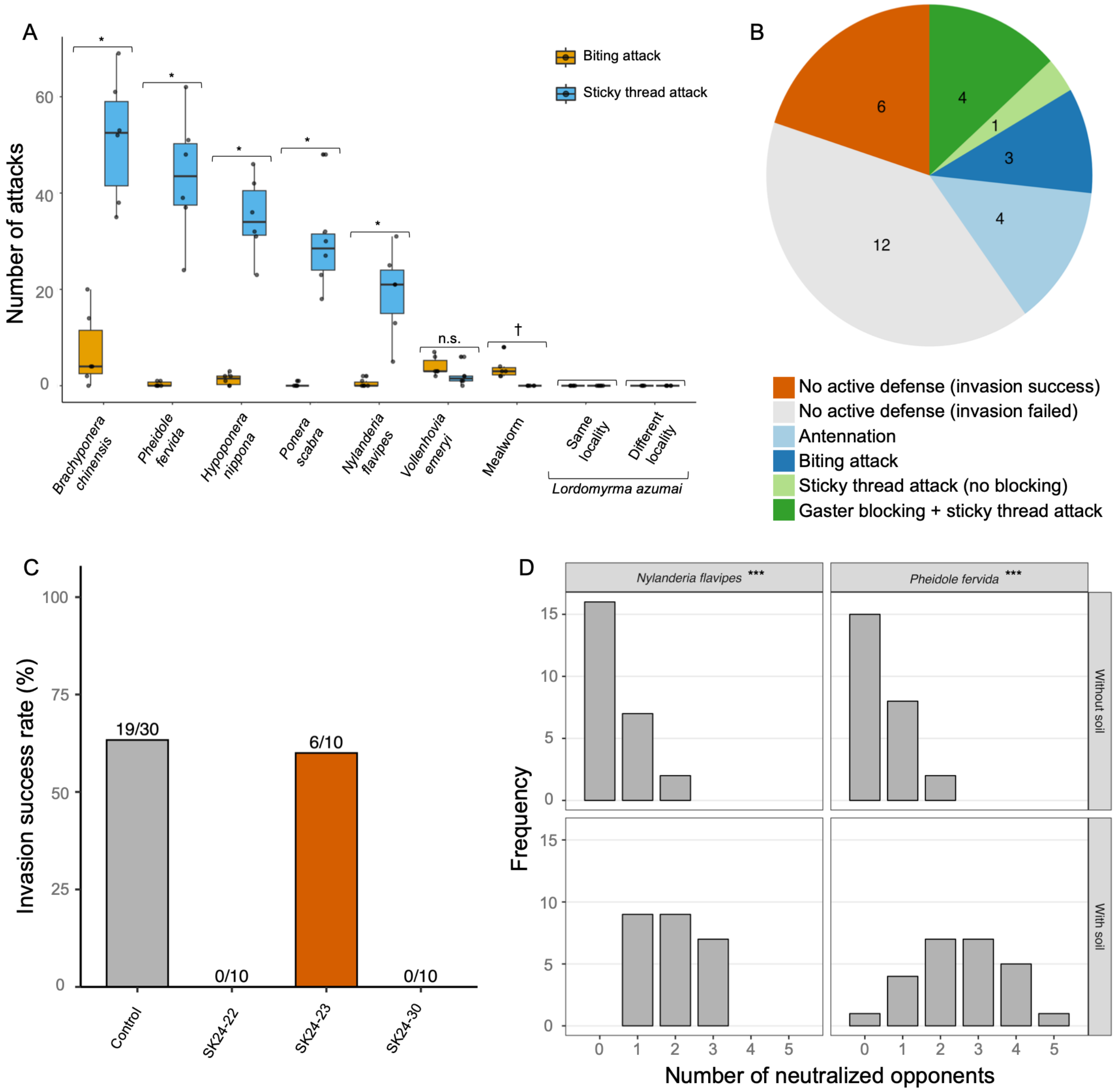


Figure S1. Context-dependent deployment of the adhesive thread and its environmentally-assisted immobilization.

(A) Frequency of defensive behaviors (biting attack versus sticky thread attack) by *L. azumai* workers against various opponents (per-opponent paired $t$-tests with colony as the unit of replication, n = 3 colonies; * Benjamini-Hochberg-adjusted $p < 0.05$; n.s., not significant). Biting attacks are shown in yellow, and sticky thread attacks in blue. No statistical comparison was performed for mealworm (†), as sticky thread attacks were absent in all replicates. No symbol is shown for conspecific opponents (*L. azumai*, same and different locality), as no aggressive behavior of any kind was observed. Box plots show median (center line), interquartile range (IQR) (box limits), and 1.5× IQR (whiskers). Individual points represent the number of attacks in each trial. Six experimental trials were performed for each opponent species. (B) Behavioral outcomes of resident workers at the nest entrance against *Nylanderia flavipes* (n = 30 trials); numbers are counts of trials in each category. All six successful invasions occurred without active defense, whereas none of the twelve trials with active defense resulted in invasion. (C) Per-colony invasion success rate of *N. flavipes* in empty control chambers versus each of three *L. azumai* colonies. (D) Frequency distribution of neutralized opponents by *L. azumai* in the presence or absence of soil. Histograms show the distribution of trial outcomes, expressed as the number of introduced opponents neutralized (0-5 of 5 individuals). The left column shows results for *N. flavipes* and the right column for *Pheidole fervida*. The top row indicates conditions without soil, and the bottom row indicates conditions with soil. The presence of soil significantly shifted the distribution toward higher neutralization numbers (Generalized linear mixed model; *** $p < 0.001$). Twenty-five experimental trials were performed for each group.

### Supplemental Experimental Procedures

**Ant colony collection and laboratory rearing.** A total of twenty-two *Lordomyrma azumai* colonies were collected between October 2024 and May 2026 in Japan. Nineteen colonies were collected from Nishiueta Town, Takamatsu City, Kagawa Prefecture. The remaining three colonies were collected from Ehime Prefecture: one from Cape Sada (Ikata Town) and two from Mount Takanawa (Matsuyama City). Colonies were maintained in custom laboratory setups consisting of a plastic brood chamber (3.5 cm long x 3.5 cm wide x 1.5 cm high) connected to a foraging arena (10 cm long x 6.5 cm wide x 2.5 cm high). The floor of the brood chamber was covered with plaster mixed with activated charcoal powder to maintain humidity. All colonies were housed under controlled laboratory conditions at 23-26°C and were regularly provisioned with mealworms (*Tenebrio molitor*). All specimens are deposited at the Laboratory of Entomology, Faculty of Agriculture, Kagawa University.

**Behavioral assays.** All behavioral observations were conducted under an Olympus SZ61 stereo microscope (Olympus, Tokyo, Japan).

**Experiment 1: Defense by single workers.** To observe the defensive responses of isolated individuals, a single *Lordomyrma azumai* foraging worker was placed in a custom-made observation container. The container consisted of a groove (3.5 cm long x 0.5 cm wide x 0.2 cm deep) carved into a plaster floor and covered with a glass plate. A single opponent ant (*Nylanderia flavipes* worker or *Pheidole fervida* minor worker) was then introduced. The frequencies of four distinct defensive behaviors–biting, sticky thread attacks, defensive posture (retracting antennae and remaining motionless), and escaping–were recorded over 30 min (n = 10 replicates per opponent species). The *L. azumai* workers used against *N. flavipes* were sourced from colonies SK24-22, SK24-25, SK25-19, and SK25-39, while those used against *P. fervida* were sourced from colonies SK25-40, SK25-41, and SK25-42. These assays were conducted between July 14 and September 27, 2025.

**Experiment 2: Defensive behavior inside the nest.** Individual opponents were introduced one at a time into the brood chambers of three *L. azumai* colonies (SK24-22, SK24-30, and

SK24-31). The entrance was plugged to prevent escape. To assess context-dependent responses, opponents were selected from three categories: (i) six sympatric ant species (*Brachyponera chinensis*, *Hypoponera nippona*, *Nylanderia flavipes*, *Pheidole fervida*, *Ponera scabra*, *Vollenhovia emeryi*); (ii) non-nestmate conspecifics, including same-locality (colony SK24-29) and different-locality (colony SK25-19) individuals; and (iii) prey items (mealworms). The frequencies of biting and sticky thread attacks were counted over 30 min (n = 6 replicates per opponent type). Assays were conducted between November 19, 2024, and July 1, 2025.

**Experiment 3: Nest entrance defense.** To evaluate defense efficiency at the nest entrance, we compared the invasion success rate of *N. flavipes* workers into *L. azumai* brood chambers (colonies SK24-22, SK24-23, and SK24-30; n = 30 trials) against empty control containers (n = 30 trials). A V-shaped partition made of Kent paper (side length: 1.5 cm, height: 1.0 cm) was placed in front of the entrance to funnel the invading opponent directly toward the entrance. We recorded the outcome (successful or failed invasion) and the specific defensive behaviors exhibited by *L. azumai* workers, classifying them into gaster blocking with thread secretion, thread attack without blocking, biting, antennation, or no active defense. Trials were conducted on March 6-7, 2025.

**Experiment 4: Nest defense effectiveness with or without soil.** We assessed the influence of soil substrate on the thread's immobilization efficiency using a crossover design for two opponent species (*N. flavipes* and *P. fervida*). Five opponents of a single species were simultaneously introduced into brood chambers with either a bare plaster floor (colonies SK25-40, SK25-43, SK25-44) or a plaster floor covered with added soil substrate (sieved to ≤ 0.7 mm; colonies SK25-41, SK25-42). Colonies were acclimated for 12 hours before the trials. After 30 minutes, the number of neutralized (i.e., immobilized and tethered to the substrate) opponents was recorded (n = 25 trials per condition). The substrate conditions for all colonies were subsequently reversed, after an interval of at least 48 hours, and the experiment was repeated. These assays were conducted between October 20 and October 29, 2025. To assess long-term lethality under soil conditions, fifteen *N. flavipes* workers were confined in the brood chambers for 30 minutes. After removing the plugs and any mobile intruders, the remaining nine immobilized *N. flavipes* were monitored at 1, 3, 5, and 24 hours on January 20-21, 2026.

**Abdominal dissection, micro-CT scanning, and scanning electron microscopy.** To identify the source of the thread secretion, we performed direct dissections. The gasters of *L. azumai* workers were dissected (n = 20 workers from two colonies) in water using fine forceps under a stereo microscope (Olympus SZ61, Japan) to examine the internal morphology and assess the physical consistency of the secretion.

The specimen for micro-CT scanning was stained in an alcoholic iodine solution for a week, followed by a 24-hour wash in 99% ethanol, and mounted in a pipette tip for scanning. Images were captured with a Zeiss Xradia 510 Versa 3D X-ray microscope (Oberkochen, Germany) operated with Zeiss Scout-and-Scan control system software (version 14.0.14829.38124) at the Okinawa Institute of Science and Technology Graduate University, Japan. The scanning parameters consisted of a 50 kV (60 μA) / 3 W beam, with a 3.5 s exposure time under 4x magnification and 2001 projections, resulting in a pixel image size of 1.37 μm. The distance between the sample and source was set to 12 mm, and the distance to the detector was 47 mm. Postprocessing of raw data was performed using ITK-SNAP 4.4.0[S1].

For the microscale structural observation of the adhesive threads, the gaster-extruded secretions were mounted on aluminum stubs using carbon tape and examined with a JEOL JCM-7000 scanning electron microscope (Tokyo, Japan) at an accelerating voltage of 15 kV.

The width of the adhesive thread was measured at 14 points along a single thread from the obtained SEM images using ImageJ software.

**FTIR spectroscopy.** To assess whether the adhesive thread is proteinaceous, we recorded Fourier-transform infrared (FTIR) spectra. Secreted threads (0.028 mg) were collected and weighed on a precision microbalance (A&D BM-20, Tokyo, Japan), mixed with 11.236 mg of potassium bromide (KBr) and pressed into a pellet; a pellet of pure KBr was used as the background blank. Spectra were acquired on a JASCO FT/IR-670 Plus spectrometer controlled by Spectra Manager for Windows 95/NT version 1.53.00, over 400–4000 $cm^{-1}$ at a resolution of 4 $cm^{-1}$ with 64 scans. The sample compartment was not purged and no atmospheric correction was applied, so water-vapor bands are present in the 1400–1800 and 3400–4000 $cm^{-1}$ regions. We therefore report only the approximate positions of the broad Amide I and Amide II envelopes; narrow features superimposed on them coincide with water-vapor lines and were not interpreted, and Amide I component bands were not resolved. The full infrared spectra are available at Zenodo[S2].

**Statistical analysis.** All statistical analyses were performed using R version 4.2.2[S3]. In Experiment 1, we compared the frequencies of each defensive behavior (biting attacks, sticky thread attacks, defensive postures, and escape) between the two opponent species (*N. flavipes* and *P. fervida*) using the Wilcoxon rank-sum test. In Experiment 2, we compared the distributions of two types of aggressive behaviors, biting attacks and sticky thread attacks, exhibited by *L. azumai* against each opponent. Because each of the three colonies (SK24-22, SK24-30, SK24-31) contributed exactly two replicates per opponent type, trials were not independent within a colony. We therefore used the colony as the unit of replication: for each opponent species, we computed the per-colony mean numbers of biting and sticky-thread attacks and compared the two attack types using a paired $t$-test ($n = 3$ colonies). $p$-values across the six opponent species were adjusted for multiple comparisons using the Benjamini-Hochberg procedure. Because only three colonies of this rare species were available at the time of this experiment, this analysis emphasizes the consistent, large-magnitude difference in attack frequency rather than precise effect-size estimation; the small number of colonies precludes formal testing of the normality assumption. In Experiment 3, successful invasions of *L. azumai* brood chambers were concentrated in a single colony; we therefore did not perform a formal statistical test for this experiment and instead report the per-colony counts of successful versus failed brood chamber invasions by opponent *N. flavipes* between the *L. azumai* colony group (n = 30 trials across 3 colonies) and the control group (n = 30 trials). In Experiment 4, we evaluated the effect of substrate presence (soil versus no soil) on the number of neutralized individuals for each opponent species. We analyzed the data using a Generalized Linear Mixed Model (GLMM) with a binomial error distribution. Substrate presence was treated as a fixed effect, and colony identity was included as a random intercept. The significance of the fixed effect was evaluated using a Likelihood Ratio Test. Model diagnostics, including tests for overdispersion, were performed using the *DHARMa* package[S4], which confirmed that overdispersion was not significant in these models ($p > 0.05$ for both opponent species). For all analyses, significance was defined as $p < 0.05$.

### Author Contributions

Conceptualization, H. Sasaki; data curation, H. Sasaki; formal analysis, H. Sasaki; funding acquisition, H. Sasaki; investigation, H. Sasaki, R.M., K.S., H. Shinohara, and K.K.; resources, H. Sasaki and F.I.; supervision, F.I.; writing – original draft, H. Sasaki; writing – review & editing, R.M., A.K., and F.I.
